# Species-specific WNT requirements shape somite morphogenesis in mammalian embryo models

**DOI:** 10.64898/2026.11.03.763751

**Authors:** Charlotte Knight, Luke Simpson, Cinzia Allegrucci, Catherine L.R. Merry, Ramiro Alberio

## Abstract

Somitogenesis establishes the metameric body plan of the vertebrate embryo and can be recapitulated *in vitro* using stem cell-based embryo models. Although these systems are well established in individual species, they have rarely been compared across species, leaving unresolved how conserved developmental programmes generate species-specific tissue organisation. Here, we compare somitogenesis in embryo models derived from human, pig, sheep and cow pluripotent stem cells and investigate how WNT signalling and the extracellular matrix regulate axis segmentation. Under identical induction conditions, human and pig models diverge in cell-type composition and morphology, revealing distinct modes of axial elongation. Increasing WNT stimulation restores somite formation in pig models but fails to restore ordered segmentation. Furthermore, each of the four mammalian species requires a different level of WNT stimulation and matrix embedding time for somite formation. Finally, comparative analyses identify species-specific differences in progenitor dynamics. Human models, showing ordered segmentation, retain a posterior mesoderm-producing domain, whereas pig models show rapid depletion of this domain, associated with simultaneous somite formation. Together, these findings reveal species-specific requirements for mammalian somitogenesis in stem cell-derived embryo models and suggest that maintenance of a posterior axial domain contributes to ordered segmentation.

## Introduction

During early development, somitogenesis establishes the segmental organisation of the vertebrate body. Shortly after gastrulation, the presomitic mesoderm (PSM) is progressively divided into bilateral somites, transient epithelial structures that give rise to the vertebrae, ribs, skeletal muscle, and dermis (Miao and Pourquié, 2024). Somites emerge in an anterior-to-posterior (AP) sequence as the body axis elongates. The position and timing are controlled by the interaction between oscillatory gene expression in the PSM and opposing gradients along the AP axis, described by the clock-and-wavefront model (Cooke and Zeeman, 1976; Hubaud and Pourquié, 2014). The tempo of this process differs markedly among species and scales with the duration of embryogenesis, with segmentation periods of approximately 2 h in mouse, 3.7 h in pig and 5 h in human embryos (Conrad et al., 2023; Lázaro et al., 2023; Matsuda et al., 2020).

Studies in vertebrate embryos show that somites arise through two temporally distinct routes. The early, anterior-most somites derive from cells that ingress through the primitive streak and contribute directly to the PSM during gastrulation. The somites of the trunk and tail originate from neuromesodermal progenitors (NMPs), a bipotent cell population that continuously generates PSM alongside the neural tube during axial elongation (Guibentif et al., 2021; Guillot et al., 2021; Wymeersch et al., 2016). As cells migrate away from the tailbud, progressive loss of *WNT3A* and *FGF8* transcripts establishes a WNT/FGF signalling gradient along the AP axis (Dubrulle and Pourquié, 2004). This gradient is opposed by anterior retinoic acid (RA) activity, positioning the determination front at which cells acquire competence for segmentation (del Corral et al., 2003). In this context, WNT signalling regulates NMP maintenance and PSM production (Garriock et al., 2015; Koch et al., 2017). It also governs the speed of posterior axis growth in embryos and stem cell-derived embryo models (Oginuma et al., 2017; Turner et al., 2017). In parallel, extensive extracellular matrix (ECM) remodelling drives the mesenchymal- to-epithelial transition that converts the PSM into morphologically distinct somites (Rifes and Thorsteinsdóttir, 2012).

Pluripotent stem cells (PSCs) are a valuable tool for studying early development *in vitro*. Under defined conditions, PSCs self-organise into embryo-like structures that recapitulate key features of gastrulation and axis formation (Arias et al., 2022). Exposure to the WNT agonist CHIR99021 (CHIR) promotes posterior patterning and elongation, while subsequent ECM embedding in Matrigel (MG) supports the emergence of more complex tissue architecture, including the formation of somites, a neural tube, or both, termed trunk-like structures (TLSs) (Girgin et al., 2021; Veenvliet et al., 2020; Yamanaka et al., 2023). Additionally, CHIR acts as a dial between neural and somitic fate, with the optimal concentration for elongation and somitogenesis typically requiring line-specific optimisation (Makwana et al., 2025). ECM embedding is, in turn, required for epithelialisation but dispensable for somitic identity, indicating that cell fate and tissue morphology can be uncoupled (Sanaki-Matsumiya et al., 2022).

The transcriptional programme of somitogenesis has predominantly been characterised in classical vertebrate models (mouse, chick and zebrafish) with some transcriptomic analyses extending to human and primate (Xi et al., 2017; Zhai et al., 2022). While the core programme and signalling pathways are conserved across vertebrates, individual species exhibit distinct developmental timing, signalling responses and embryo morphologies (Ramesh and Chu, 2024; Xi et al., 2017; Zhai et al., 2022). Whether these differences are reflected in stem cell-derived models remains unclear. PSC-derived models of somitogenesis have mainly been established in mouse and human PSCs (Veenvliet et al., 2020; Yamanaka et al., 2023). In a recent study, TLSs were generated from sheep PSCs, but pig PSCs failed to form these structures under the same protocol, and the reason for this remains unclear (Hauser et al., 2026). In the pig, somites start forming around embryonic day (E) 14 (Simpson et al., 2024). Two recent studies have characterised gastrulation and cell commitment within somite derivatives (E11.5 to E15 and E16 to E28, respectively), yet the molecular underpinnings of somitogenesis remain unexplored (Cai et al., 2023; Simpson et al., 2024). Direct comparisons between species remain challenging because PSC maintenance conditions influence developmental competence, signalling responsiveness and lineage composition in gastruloids (Anlas et al., 2025; Balayo et al., 2025). Consequently, differences observed between published models may reflect variation in starting cell state rather than genuine biological divergence. A systematic comparison using common culture and differentiation conditions is therefore needed to distinguish conserved features of mammalian somitogenesis from species-specific developmental requirements. In this study, we examined PSM development in pig embryos and identified a conserved transcriptional programme of somitogenesis. We then recapitulated somitogenesis in vitro using PSCs from four species (human, pig, sheep and cow) maintained and differentiated under identical conditions and showed that tissue patterning depends on species-specific WNT activity and timing of ECM embedding. By defining how these signals influence tissue patterning, we provide a framework for understanding how conserved developmental programmes generate distinct morphological outcomes.

## Results

### The somitogenesis programme is conserved in the pig embryo

To determine whether the differences in TLS formation in the pig could result from embryological differences, we investigated the somitogenesis trajectory in pig embryos. We integrated two published single-cell datasets spanning contiguous developmental windows: a gastrulation atlas covering E11.5 to E15 (Simpson et al., 2024) and a myogenesis-focused dataset covering E16 to E28 (Cai et al., 2023). Together, these capture the full trajectory from gastrulation through somitogenesis to skeletal muscle differentiation. Following integration, the datasets were annotated, subset to the clusters of interest and their neighbouring populations and finally reannotated to resolve cell types at higher resolution (Fig. 1A-C). The resulting clusters showed the expected developmental sequence, progressing from epiblast and primitive streak through to nascent mesoderm (NM), PSM and somitic mesoderm, to dermomyotome and the myogenic lineages (Fig. 1A). Cluster identities were assigned using an extended marker panel for each subcluster (Suppl. Fig. 1A). The canonical stage markers delineated the clusters leading to somitogenesis, with *TBXT* marking NM, *TBX6* marking the PSM and *MEOX1* and *PAX3* marking somitic mesoderm (Fig. 1D). However, *PAX3* expression extended to the neural tube cluster, showing it cannot be used independently as a somite marker.

**Figure 1.**
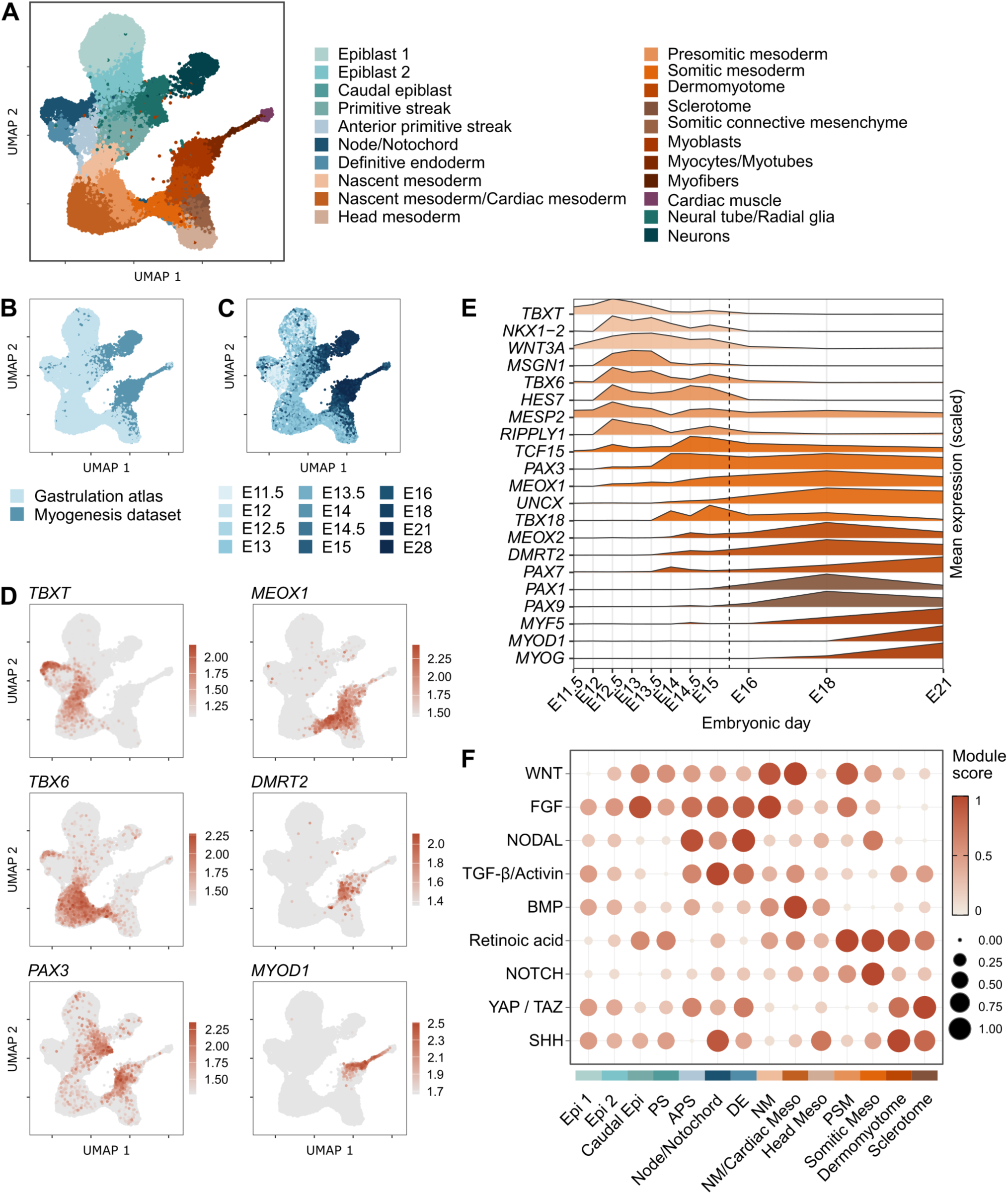
A single-cell transcriptomic atlas of pig somitogenesis. (A) UMAP of integrated single-cell datasets from pig embryos between E11.5 and E28, subset to the somitogenesis trajectory and neighbouring populations. Cells are coloured by annotated cell type. (B) UMAP coloured by dataset of origin: gastrulation atlas (E11.5-E15; Simpson et al., 2024) and myogenesis dataset (E16-E28; Cai et al., 2023). (C) UMAP coloured by embryonic day. (D) Feature plots showing the expression of TBXT, TBX6, PAX3, MEOX1, DMRT2 and MYOD1. Cells are coloured by log-normalised expression (RNA assay). (E) Mean log-normalised expression of marker genes across embryonic days (E11.5-E21), scaled to each gene’s maximum. Genes are coloured by population: posterior progenitors (*TBXT, NKX1-2, WNT3A*), presomitic mesoderm (*MSGN1, TBX6, HES7, MESP2, RIPPLY1*), somite (*TCF15, PAX3, MEOX1, UNCX, TBX18*), dermomyotome (*MEOX2, DMRT2, PAX7*), sclerotome (*PAX1, PAX9*) and myogenic cells (*MYF5, MYOD1, MYOG*). The dashed line marks the boundary between the two datasets. (F) Dot plot of signalling pathway module scores across the 14 clusters covering the somitogenesis trajectory. Dot colour and size both represent the scaled module score. Epi, epiblast; PS, primitive streak; APS, anterior primitive streak; DE, definitive endoderm; NM, nascent mesoderm; Meso, mesoderm; PSM, presomitic mesoderm.

To resolve the timeline of lineage progression from gastrulation to myogenic differentiation, we examined marker gene expression and the relative abundance of key cell populations across embryonic stages (Fig. 1E, Suppl. Fig. 1B). Expression of the NMP markers *TBXT*, *NKX1-2* and *WNT3A* peaked at E12.5, coinciding with the onset of secondary gastrulation following the formation of the first notochord cells. The PSM regulators *MSGN1*, *TBX6*, *HES7*, *MESP2* and *RIPPLY1* also peaked in expression during this period. Notably, the somitic mesoderm markers *PAX3* and *TCF15* as well as the rostral somite marker *TBX18* increased rapidly between E13.5 and E14.5, around the formation of the first six somites (Simpson et al., 2024). Dermomyotome cells were first detectable at E14, concomitant with increased expression of the dermomyotome markers *MEOX2* and *DMRT2*. The dermomyotome cluster appeared prior to the *PAX1*- and *PAX9*-expressing sclerotome, which was only detected from E15 onwards. This suggests that by the formation of the sixth somite, the anterior somites have already acquired rostro-caudal identity and at least some have undergone dorsolateral patterning. Interestingly, this appears to precede ventromedial patterning by one day. Myogenic regulatory factors were induced sequentially, with *MYF5* increasing at E18 followed by *MYOD1* and *MYOG* at E21, marking the formation of myoblasts and myocytes, respectively.

We then asked whether the signalling dynamics underlying this trajectory were conserved. We grouped ligands, receptors and transcriptional targets into pathway-specific modules (Suppl. Table 1) and scored these across the annotated clusters. Pig embryos recapitulated the spatiotemporal signalling patterns described in primate embryos (Xi et al., 2017; Zhai et al., 2022). WNT signalling was highest in NM and PSM, FGF signalling peaked in the primitive streak and NM, and RA signalling was elevated in PSM and somitic mesoderm, consistent with the anterior RA gradient required for segmentation (Fig. 1F).

To gain spatial information on somitogenesis markers, we performed whole-mount immunostaining of 8-somite stage pig embryos. Embryos were divided into an anterior portion, containing the somites and the brain, and a posterior portion, containing the PSM and tailbud (Fig. 2A). In the anterior portion, PAX3 marked the somites that flank the neural tube, which was positive for SOX1 and SOX2 (Fig. 2B, C). In the posterior portion, SOX2 and TBXT were co-expressed in a discrete domain in the tailbud, corresponding to the NMP population (Fig. 2E). Immediately anterior to this domain, the NMPs fed into two distinct structures: a SOX2-positive neural tube and a TBX6-positive PSM occupying a narrow band that transitioned anteriorly into unsegmented somitic mesoderm (Fig. 2D). This spatial organisation is consistent with the presence of a bipotent progenitor pool supplying both neural and mesodermal lineages during axial elongation. Together, these patterns are consistent with those described in mouse, chick, monkey and zebrafish, demonstrating that mesoderm specification and somitic lineage commitment in the pig embryo follow a similar transcriptional programme, signalling dynamics and spatial organisation.

**Figure 2.**
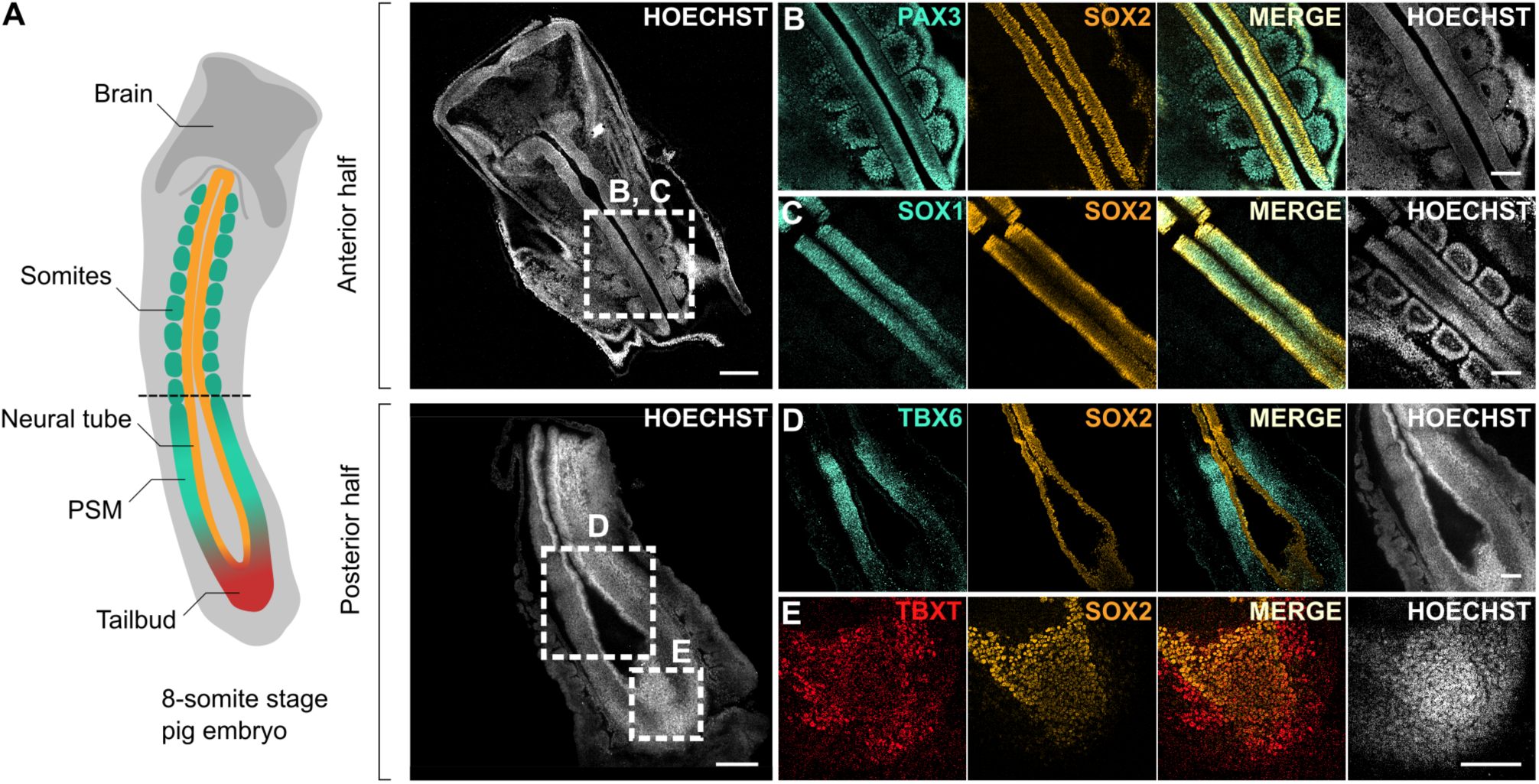
Immunofluorescence characterisation of the trunk and posterior domains in 8-somite stage pig embryos. (A) Schematic of an 8-somite stage pig embryo alongside overviews of the anterior half (top; including trunk and head) and posterior half (bottom; including the PSM region and tailbud) of two embryos. Boxes indicate the domains shown in (B-E). (B) PAX3 and SOX2 immunostaining of the trunk domain. (C) SOX1 and SOX2 immunostaining of the trunk domain in a same-stage embryo. (D) TBX6 and SOX2 immunostaining of the PSM domain. (E) SOX2 and TBXT immunostaining of the NMP domain. Embryos were immunostained as whole-mounts; all images show a single z-plane. Three stage-matched embryos were used, shown in (B), (C) and (D,E) respectively. Scale bars: 200 µm (A), 100 µm (B-E).

### Pig and human embryo models diverge in fate and morphology under identical induction conditions

Having established that the pig embryo uses a conserved somitogenesis programme *in vivo*, we asked whether pig PSCs execute this programme under conditions established for human PSCs. Using an established protocol (Yamanaka et al., 2023), we compared two human (H9 and HUES7) and two pig (pEpiSC7 and pEpiSC4, hereafter P7 and P4) PSC lines, all maintained under identical culture conditions (see methods). Gastruloids were embedded in MG at two different timepoints and CHIR was tested across a range of 0.5 to 5 µM to identify the optimum for each species (Fig. 3A). We refer to these aggregates as gastruloids prior to MG embedding, and as axioloids or TLSs after embedding, once higher-order tissue organisation emerges. Neither species elongated at the lowest CHIR concentrations. Human gastruloids elongated robustly at intermediate doses (Fig. 3B, C), forming axioloids with sequential MEOX1-positive somites by 96 h following MG embedding at 72 h (Fig. 3D). At the highest dose, human gastruloids either did not elongate or formed multiple poles with reduced elongation. Upon embedding in MG, these produced fewer and smaller somites than at the intermediate dose (Fig. 3B). In contrast, pig gastruloids elongated at both intermediate and high doses, with the most pronounced elongation at the highest dose (Fig. 3B, C). Unlike human, pig gastruloids were already visibly elongating by 24 h under this regime. Aspect ratio (AR) measurements revealed differences between species across CHIR conditions and timepoints, with pig AR increasing earlier than human but collapsing by 96 h across all conditions (Fig. 3C). Despite this capacity to elongate, pig gastruloids did not form somites at either CHIR dose or embedding timepoint. Following embedding at 48 h, pig gastruloids at 72 h were predominantly SOX2-positive, containing only a small anterior MEOX1-positive domain (Fig. 3D). Human axioloids contained a discrete SOX2/TBXT double-positive population restricted to the posterior tip, corresponding to NMPs. In pig, TBXT was present but marked a central region within the gastruloid, more consistent with axial mesoderm/notochord identity (Fig. 3D). Equivalent morphologies were observed in other human and pig lines, indicating that the phenotype was not cell line-specific (Suppl. Fig. 2A, B).

**Figure 3.**
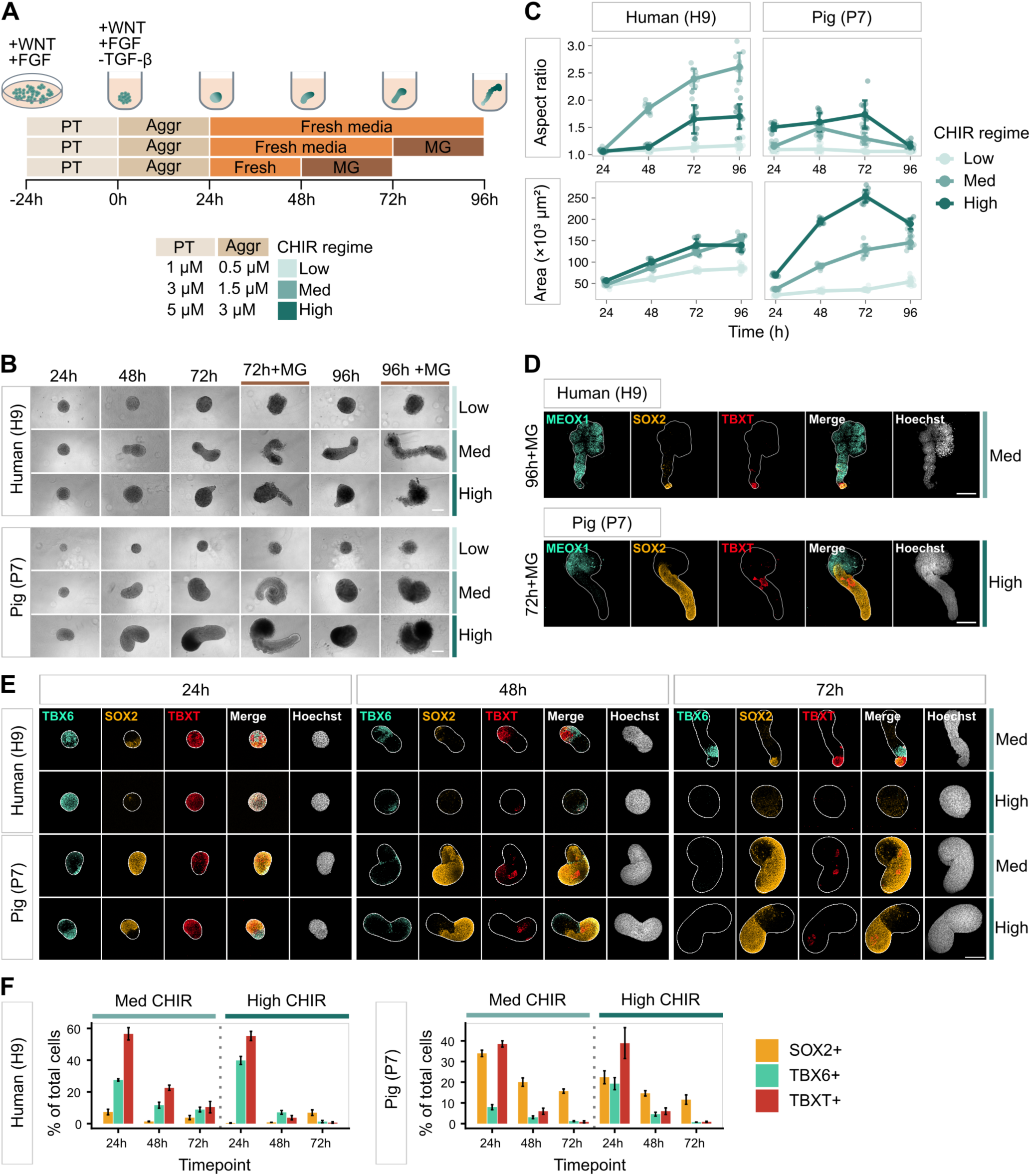
Pig and human gastruloids diverge in fate and morphology under identical conditions. (A) Schematic of the gastruloid protocol. Cells were pre-treated (PT) for 24 h with FGF and CHIR and aggregated (Aggr) with FGF, CHIR and SB431542. Gastruloids were maintained in fresh medium thereafter and embedded in Matrigel (MG) at either 48 or 72 h. CHIR regimes (low, med, high) are defined in the table. (B) Brightfield images of human (H9) and pig (P7) gastruloids from 24 to 96 h under low, med and high CHIR regimes. 72 h + MG and 96 h + MG indicate gastruloids embedded in MG 24 h before imaging. Scale bar, 200 µm. (C) Aspect ratio (top) and area (bottom) of human (H9) and pig (P7) gastruloids over time under the three CHIR regimes. Data show mean ± s.d., with points representing individual gastruloids (n = 9-12 gastruloids per condition). (D) Immunofluorescence images (maximum-intensity projections) of human (H9, med CHIR, 96 h + MG) and pig (P7, high CHIR, 72 h + MG) gastruloids stained for MEOX1, SOX2 and TBXT, with Hoechst. White lines indicate the gastruloid outline. Scale bars, 200 µm. (E) Immunofluorescence images (maximum-intensity projections) of human (H9) and pig (P7) gastruloids at 24, 48 and 72 h under med and high CHIR regimes, stained for TBX6, SOX2 and TBXT, with Hoechst. White lines indicate the gastruloid outline. Scale bar, 200 µm. (F) Percentage of total cells positive for SOX2, TBX6 and TBXT in human (H9) and pig (P7) gastruloids at 24, 48 and 72 h under med and high CHIR regimes. Marker-positive populations are not mutually exclusive. Bars show mean ± s.d. (n = 3 gastruloids per condition).

We next examined marker dynamics over the first 72 h of the protocol to determine when the two species diverged. We stained for TBX6, SOX2 and TBXT at 24, 48 and 72 h under the conditions permissive for elongation in each species. In human with intermediate CHIR, a substantial TBX6-positive population was established by 24 h and persisted through to 72 h, progressively restricting to a domain immediately anterior to the SOX2/TBXT-positive tailbud (Fig. 3E). At the highest CHIR dose, this organisation was lost. TBX6 was induced throughout the aggregate by 24 h and largely lost by 48 h, with little SOX2 or TBXT remaining, indicating that no posterior PSM or NMP population was retained. Raising WNT therefore accelerated differentiation at the expense of the progenitor pool, and with it the morphology that depended on continued progenitor output.

In pig with high CHIR, TBX6 was expressed at 24 h, then was nearly absent by 48 h and undetectable by 72 h. At the intermediate dose, a substantial TBX6-positive population was never established. In both conditions, SOX2 expanded over time to dominate the structure. Quantification of marker expression confirmed these opposing trends (Fig. 3F). SOX2/TBXT double-positive cells were abundant at 24 h but were hardly detectable by 48 h. At this stage, TBXT marked a central population lying outside the tailbud, consistent with axial mesoderm identity and previous reports of notochord in pig gastruloids (Hauser et al., 2026). NMPs were therefore initially specified in pig gastruloids but did not sustain a PSM compartment, leaving no progenitor supply to feed a somitic population and no segmentation.

These data showed that pig gastruloids presented an unexpected phenotype under standard induction conditions. At the CHIR concentration that abolished correct elongation in human, pig gastruloid elongation was the greatest, starting at 24 h, yet the resulting structures were predominantly neural and did not segment. Pig aggregates remained as gastruloids under these conditions, without acquiring higher-order organisation. Elongation in the two species proceeded through different mechanisms and on different timescales: in human through mesoderm differentiating from NMPs in the tailbud, and in pig through expansion of neural tissue. Therefore, although elongation was required for proper somite formation, it was not sufficient. The maintenance of a pool of mesodermal progenitors was also required.

### Increased CHIR overcomes reduced WNT sensitivity but does not restore somite organisation in pig TLSs

To establish whether pig and human cells differed in their sensitivity to CHIR dose independently of 3D self-organisation, we exposed monolayer cultures of both species to a two-day CHIR regime ranging from 0.5 to 15 µM and quantified the proportion of TBX6- and SOX2-positive cells in each condition (Fig. 4A). The two species showed distinct patterns: the peak in TBX6-positive fractions occurred at the higher end of the CHIR range in pig but at intermediate doses in human. On the other hand, SOX2 was retained at higher doses in pig than in human, where the proportion of SOX2-positive cells declined sharply as CHIR dose increased (Fig. 4B). The dose-fate relationship was therefore shifted in pig cells, which required more CHIR than human to reach a comparable balance of TBX6- and SOX2-positive cells. Human cells died at the highest dose and the wells contained too few cells for reliable quantification (Suppl. Fig. 3A). The two species therefore differed in CHIR sensitivity even without the morphogenetic context of a self-organising aggregate.

**Figure 4.**
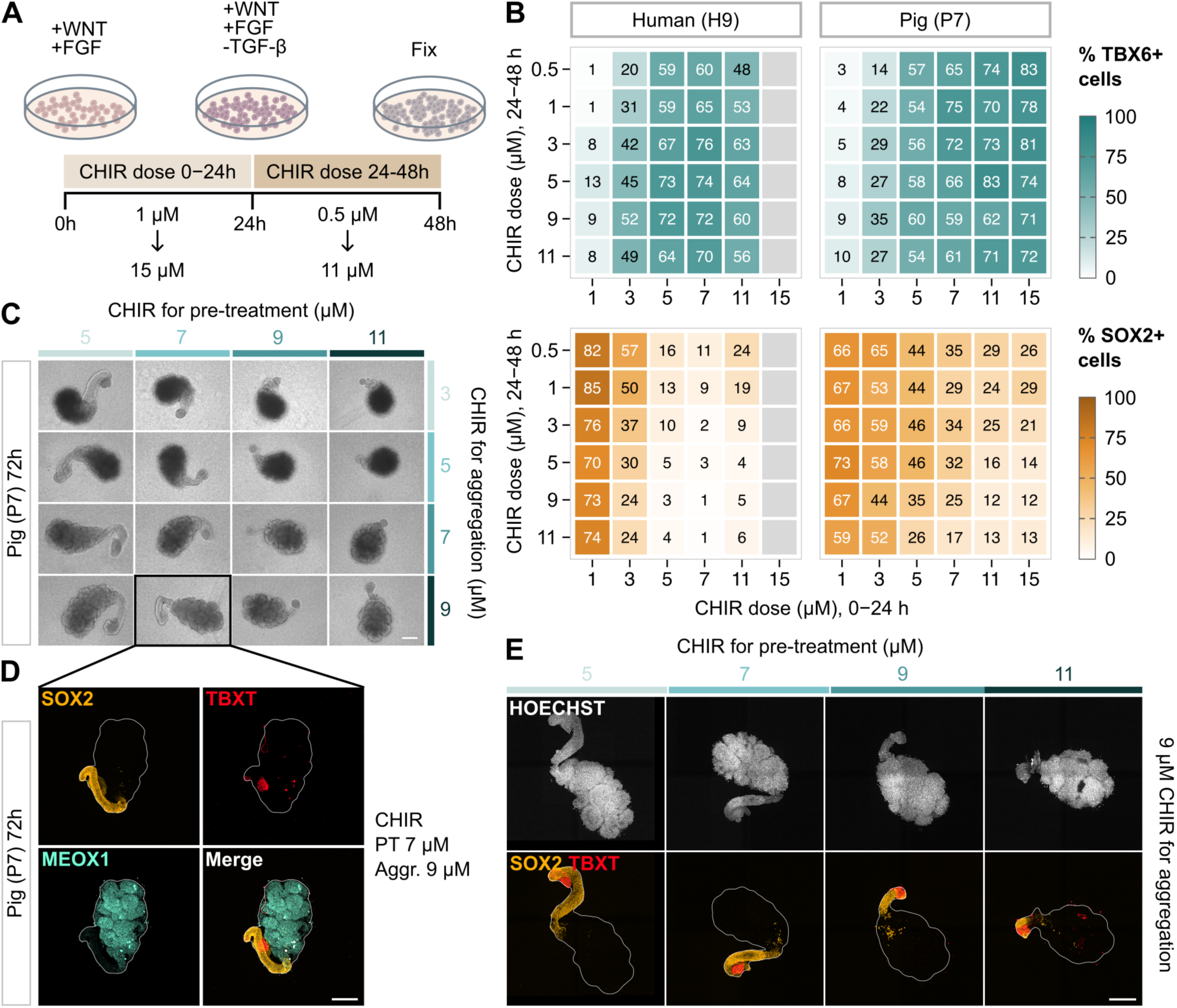
Pig gastruloids require higher CHIR doses for somite formation. (A) Schematic of the 2D CHIR dose-response experiment. Cells were exposed to CHIR from 0 to 24 h (1-15 µM) and from 24 to 48 h (0.5-11 µM) in the presence of FGF, with SB431542 from 24-48 h, and were fixed at 48 h. (B) Percentage of TBX6+ (teal) and SOX2+ (orange) cells in human (H9) and pig (P7) across a matrix of CHIR doses. The x-axis shows the CHIR dose at 0-24 h and the y-axis the dose at 24-48 h. Each value is the mean of 3 regions per well across 3 independent experiments. Grey squares indicate conditions excluded from quantification because of insufficient cell numbers. (C) Brightfield images of pig (P7) gastruloids at 72 h + MG across a matrix of CHIR concentrations at pre-treatment (columns) and aggregation (rows). The box marks the condition immunostained in D (pre-treatment 7 µM, aggregation 9 µM). (D) Immunofluorescence images (maximum-intensity projections) of a pig (P7) TLS at 72 h + MG stained for SOX2, TBXT and MEOX1. The white line indicates the TLS outline. (E) Immunofluorescence images (maximum-intensity projections) of pig (P7) TLSs at 72 h + MG generated with different CHIR concentrations at pre-treatment and 9 µM CHIR at aggregation. Top: Hoechst. Bottom: merge of SOX2 and TBXT. White lines indicate the TLS outline. Scale bars, 200 µm.

Given the differential sensitivity and the tolerance of pig cells to such high CHIR concentrations, we hypothesised that raising the level of WNT signalling in pig gastruloids would shift the balance between neural and mesodermal fates. We tested CHIR doses above the standard range to determine whether this was sufficient to generate the mesodermal progenitors required for somite formation, without compromising elongation. In the following experiments, gastruloids were embedded in MG 24 h before each timepoint shown. Across an extended range of CHIR pre-treatment and aggregation conditions, pig gastruloids still elongated at CHIR concentrations several-fold above the standard concentrations. At the highest concentrations (upwards of 5 and 7 µM for pre-treatment and aggregation, respectively), they formed TLSs at 72 h showing a “bunch of grapes” phenotype (Dias et al., 2014; Veenvliet et al., 2020) (Fig. 4C, Suppl. Fig. 3B). These clusters were MEOX1-positive, confirming somitic identity. Surprisingly, TLSs retained a SOX2-positive neural tube domain posteriorly at high CHIR concentrations (Fig. 4D). The SOX2-positive domain contracted progressively as CHIR dose increased (Fig. 4E). Increasing the levels of WNT signalling therefore shifted the balance towards mesoderm and produced sufficient levels of progenitors to enable somite formation. However, this was not sufficient to rescue somite organisation.

### Species-specific requirements for WNT stimulation and ECM embedding for *in vitro* somitogenesis

Having found that pig cells require CHIR concentrations outside the range that supports somitogenesis in human cells, we asked whether species-specific CHIR and MG embedding requirements extended to other mammals. We tested a similar CHIR dose range in sheep and cow PSCs (sEpiSC21 and bEpiSC99.1, hereafter S21 and B99.1) together with two different embedding timepoints, with all lines maintained under identical culture conditions. Each cell line showed distinct optima for somitogenesis (Fig. 5A, B). Sheep gastruloids formed TLSs with organised somites at an intermediate CHIR dose, higher than the human optimum but lower than pig, and best when embedded at 48 h. Cow gastruloids instead required a higher dose, closer to the pig range, and embedding at 72 h. Interestingly, delaying embedding to 72 h in sheep still produced somites, but fewer in number, restricted to the anterior and to one side of the axis. AR measurements across four CHIR regimes confirmed that the two species elongated differently across the same conditions, with sheep reaching maximal elongation at lower doses than cow (Fig. 5C, Suppl. Fig. 4). Immunostaining confirmed that the somites were negative for TBXT and SOX2 (Fig. 5D). In both species, somites were distributed along the AP axis rather than confined to a large anterior cluster, as seen in pig, but neither species reproduced the defined sequential organisation seen in human. At doses slightly above their optimum, sheep TLSs formed structures resembling the “bunch of grapes” phenotype, although these were spread along the length of the axis. Cow TLSs formed somites along the axis but also produced disorganised clusters anteriorly. Both species retained a posterior SOX2-positive domain, larger than that of human axioloids, and in cow this domain persisted even at the highest CHIR concentrations tested.

**Figure 5.**
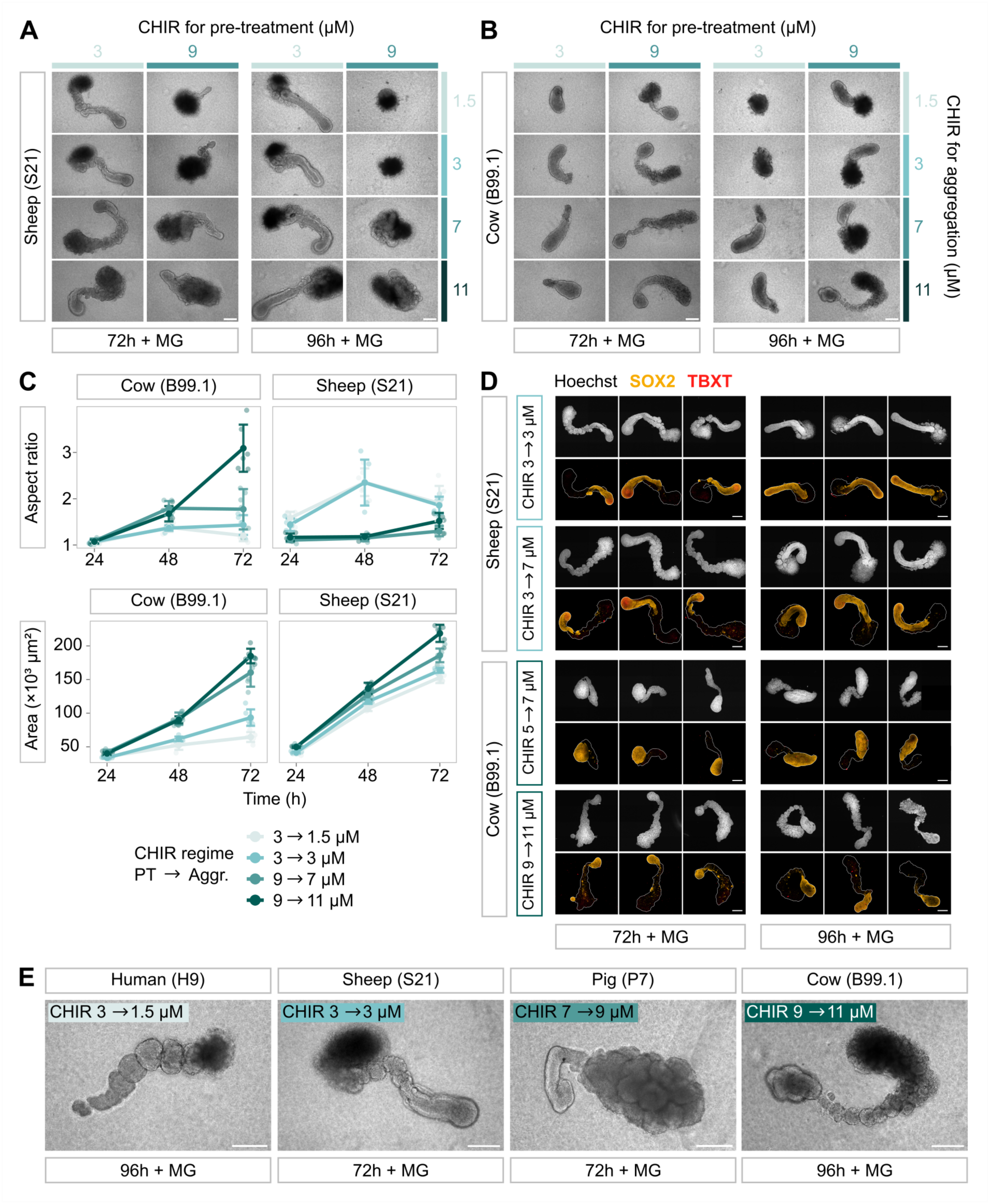
CHIR and MG embedding requirements vary across four mammalian species. (A) Brightfield images of sheep (S21) gastruloids at 72 h and 96 h (embedded in MG 24 h prior) across a matrix of CHIR concentrations at pre-treatment (columns) and aggregation (rows). (B) Same as A, for cow (B99.1) gastruloids. (C) Aspect ratio and area of cow and sheep gastruloids over time under four CHIR regimes (pre-treatment/aggregation: 3/1.5, 3/3, 9/7 and 9/11 µM). Data show mean ± s.d., with points representing individual gastruloids (n = 6 gastruloids per condition). (D) Immunofluorescence images (maximum-intensity projections) of sheep and cow TLSs at 72 h + MG and 96 h + MG stained for SOX2 and TBXT. Each species is shown under two CHIR regimes (pre-treatment/aggregation: sheep, 3/3 and 3/7 µM; cow, 5/7 and 9/11 µM). Columns show TLS replicates. Top rows: Hoechst. Bottom rows: merge of SOX2 and TBXT. White lines indicate the TLS outline. (E) Brightfield images of human, sheep, pig and cow embryo models under the condition that gave the best somite formation for each species, labelled with timepoint and CHIR regime. Scale bars, 200 µm.

Overall, across the four species tested starting from an identical pluripotent culture condition, each required a distinct CHIR dose regime and MG embedding time for somite formation (Fig. 5E). Sheep and cow TLSs showed an intermediate phenotype between human and pig, with somites distributed along the axis but without forming a single regular row. These results suggest that species differ in two aspects: the level of WNT required to produce enough PSM for somite formation, and the extent to which neural tissue contributes to elongation. This also suggests that a specific ratio of neural and mesodermal tissues is required for the correct patterning of somites. The ordering of CHIR requirements, with human lowest, followed by sheep, then cow and pig at the highest doses, showed no clear relationship with known differences in developmental tempo or phylogenetic distance between these species.

### Loss of progenitors is associated with simultaneous rather than sequential segmentation in pig TLSs

To further characterise the mechanism of segmentation in the most divergent model, we compared the timing and spatial order of somitic boundaries in human and pig models. Using CHIR pre-treatment and aggregation doses of 3 and 1.5 µM for human and 11 and 9 µM for pig, we performed live imaging of axioloids and TLSs from the point of MG embedding over a 20 h window (Suppl. Movies 1 and 2). We used the appearance of grooves to mark the emergence of somite boundaries (Fig. 6A). In human axioloids, boundaries appeared progressively over time, beginning anteriorly and extending posteriorly, recapitulating the segmentation dynamics observed *in vivo* (Fig. 6B). In pig TLSs, boundaries formed within a narrower time window (3.4 ± 1.7 h in pig vs 12.8 ± 2.3 h in human) and without consistent AP order, with the posterior half completing segmentation at the same time as the anterior half (Fig. 6B). This pattern is inconsistent with a determination front travelling along the AP axis.

**Figure 6.**
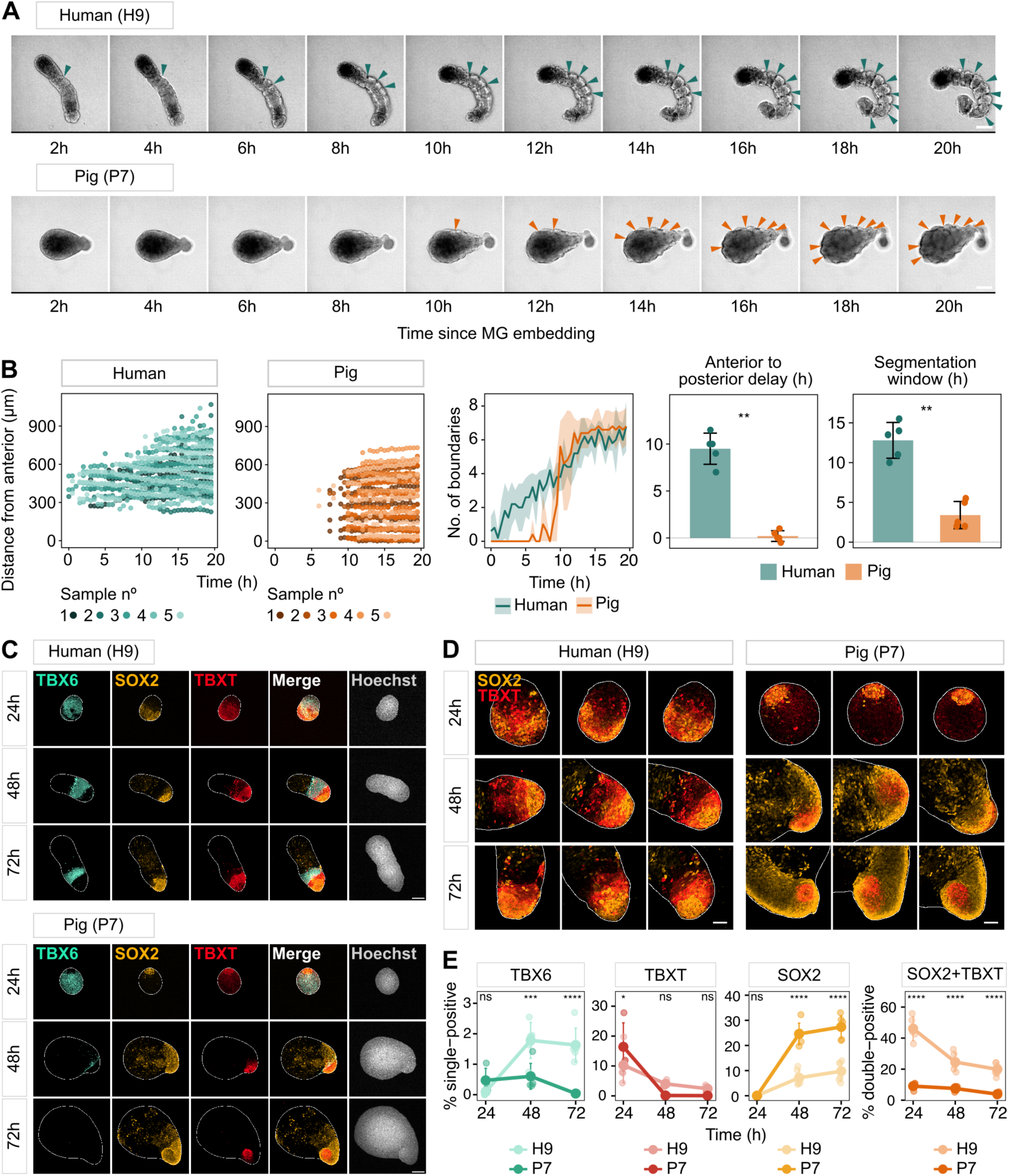
Pig and human embryo models differ in segmentation dynamics and progenitor organisation. (A) Brightfield snapshots from live imaging of human (H9) axioloids and pig (P7) TLSs (CHIR pre-treatment/aggregation: 3/1.5 µM for human, 11/9 µM for pig). Snapshots are shown every 2 h, from 2 to 20 h post MG embedding (embedded at 72 h for human and 48 h for pig). Arrows indicate somite boundaries (grooves). (B) Somite boundary dynamics in human (H9) axioloids and pig (P7) TLSs over 20 h of live imaging post MG embedding (n = 5 per species). Left: position of each boundary along the AP axis (distance from the anterior end) over time; each point is one boundary in one frame, coloured by replicate. Middle: number of boundaries over time (mean ± s.d.). Right: anterior-to-posterior delay (difference in the time taken by the posterior and anterior halves to complete segmentation) and segmentation window (time taken to form all boundaries), shown as mean ± s.d. Mann-Whitney test; **P < 0.01. (C) Immunofluorescence images (maximum-intensity projections) of human (H9) and pig (P7) gastruloids (CHIR pre-treatment/aggregation: 3/1.5 µM for human, 11/9 µM for pig) at 24, 48 and 72 h stained for TBX6, SOX2 and TBXT, with Hoechst. White lines indicate the gastruloid outline. (D) Immunofluorescence images (maximum-intensity projections) of the posterior tip of human (H9) and pig (P7) gastruloids at 24, 48 and 72 h showing SOX2 and TBXT (merged). Three gastruloids are shown per timepoint. White lines indicate the gastruloid outline. (E) Percentage of total cells positive for TBX6 only, TBXT only, SOX2 only, and both SOX2 and TBXT in human (H9) and pig (P7) gastruloids at 24, 48 and 72 h. Large points show mean ± s.d.; small points represent individual gastruloids (n = 4-6 gastruloids per timepoint). Two-way ANOVA with Šídák’s multiple comparisons; ns P ≥ 0.05, *P < 0.05, ***P < 0.001, ****P < 0.0001. Scale bars, 200 µm (A, C) and 50 µm (D).

To better understand this disorganised segmentation, we compared progenitor population dynamics leading up to this phenotype. In human gastruloids, the TBX6- positive domain was maintained throughout the time course, as observed previously (Fig. 6C, Suppl. Fig. 5). In pig, TBX6 was strongly induced at 24 h but rapidly downregulated thereafter and was replaced by a large anterior domain, which later gave rise to the somitic clusters. At this CHIR concentration, the TBXT-positive notochord-like domain was no longer present; instead, a small TBXT-positive population was present in the tailbud region of the gastruloid. Over the time course, the SOX2-positive domain expanded, despite being relatively small at 24 h.

We next examined the posterior tip of the gastruloid, imaging TBXT and SOX2 at higher magnification across the same time course. At 72 h, human gastruloids showed spatial organisation of SOX2 and TBXT consistent with embryo axial elongation, with a SOX2/TBXT double-positive population at the posterior tip and a TBXT-only mesodermal population emerging immediately anterior to it (Fig. 6D). Pig gastruloids showed the inverse pattern, with a SOX2-only population emerging anterior to the double-positive tip. Quantification of marker expression across the whole gastruloid confirmed this asymmetry, with a larger TBXT-only fraction in human gastruloids and a larger SOX2-only fraction in pig (Fig. 6E). There were also fewer double-positive cells for SOX2 and TBXT in pig gastruloids at 48 and 72 h, indicating a possible reduction in the NMP pool compared to human. To test whether prolonged WNT activation could rescue this phenotype and sustain mesodermal differentiation from NMPs, we continued CHIR treatment from 24 to 48 h (Suppl. Fig. 6A). Additional CHIR reduced elongation at 48 h and did not produce sequentially organised somites. At 72 h, at the highest pre-treatment and aggregation dose (11/9 µM), it also prevented the formation of somite clusters (Suppl. Fig. 6B, C).

In pig gastruloids, the posterior progenitor population therefore did not sustain a spatially organised supply of new PSM. TBX6 was lost by 48 h, with the whole PSM compartment converting to somitic mesoderm at once, precluding a progressively retracting determination front. Subsequently, the tailbud continued to generate cells that were predominantly neural rather than mesodermal. These observations indicate that the pig phenotype does not reflect an inability to specify somitic identity but a failure to sustain the progenitor dynamics that convert that identity into an ordered, sequentially segmenting axis. These findings suggest that sustained progenitor output may contribute to sequential segmentation, distinct from the capacity to specify somitic identity.

## Discussion

A central question in vertebrate development is how conserved gene regulatory networks generate species-specific patterns of tissue organisation during somitogenesis. Using stem cell-derived embryo models from four mammalian species, we show that identical culture conditions give rise to distinct cellular compositions and morphologies, revealing profound interspecies differences in developmental responses. Our findings demonstrate that somite formation depends on species-specific sensitivities to WNT signalling and uncover an unexpected relationship between the balance of mesodermal progenitors and somite morphology.

In the pig embryo, our *in vivo* analyses showed that the key features of vertebrate somitogenesis and axial elongation were conserved. Mesoderm specification, PSM formation and somite patterning proceeded through conserved genetic programmes and signalling dynamics. However, under the same induction conditions, pig and human gastruloids differed markedly in cell type and morphology. Pig gastruloids elongated through expansion of the neural domain whereas human gastruloids elongated through mesoderm production from NMPs. This is consistent with the observation that elongation does not require a fixed ratio of neural to mesodermal tissue (Anlas et al., 2025). A certain level of heterogeneity was nonetheless required. At the highest CHIR dose, human gastruloids induced TBX6 throughout the aggregate and did not elongate or did so with multiple poles, showing that a uniformly mesodermal aggregate could not extend normally. Furthermore, pig gastruloids elongated earlier than human. This may reflect the faster developmental tempo of the pig, but it may also be a consequence of which tissue drives extension. Neural expansion may induce axis elongation more rapidly than the production of mesoderm. This is consistent with the finding that mouse gastruloids elongating earlier are enriched for neural tissue and contain less PSM (Balayo et al., 2025).

Human, pig, sheep and cow cells each had distinct CHIR optima for somite formation, spanning concentrations several-fold above those used in published mouse and human protocols. These optima did not seem to track phylogeny. MG embedding time, in contrast, appeared to reflect how soon somitogenesis follows gastrulation *in vivo* in each species. This suggests that the embedding time must be matched to the developmental stage of the gastruloid rather than to a fixed culture duration. In pig TLSs, increased CHIR restored somitic identity, consistent with a shifted WNT sensitivity, but somites lacked the sequential order and axial organisation seen in human axioloids. Sheep and cow TLSs also formed clusters of somites at specific CHIR doses, though somites remained distributed along the length of the axis. A similar phenotype has been reported in mouse gastruloids exposed to high CHIR (Veenvliet et al., 2020) and in human segmentoids, in which the most anterior region often comprises grape-like rosettes lacking clear AP identity (Miao et al., 2023). Although sheep and cow phenotypes were reproducible within each line, the comparison across species is limited by the use of a single PSC line for these species.

To understand why segmentation failed to proceed sequentially in pig, we compared progenitor populations in the timepoints leading up to segmentation. TBX6 was induced by 24 h in both species, but in human axioloids it was maintained and progressively restricted to a domain anterior to the tailbud. In pig TLSs, TBX6 was lost by 48 h and the NMPs never replenished the mesodermal population. Similarly, in human, above the WNT optimum, the PSM was not retained and fewer, smaller somites formed. In both species, WNT levels set not only whether PSM was produced, but also whether it was maintained, and this determined how somites formed. In the embryo, somitogenesis ends when the PSM pool is exhausted (Gomez et al., 2008). Our results show the same dependency on a shorter timescale, with increased WNT spending the progenitor pool early and reducing the number of somites formed. Progenitor supply may therefore be one point of control over segment number between species.

Furthermore, the early loss of PSM in pig TLSs and the neural differentiation of NMPs suggest that somites formed via the early, NMP-independent route, with NMPs supplying SOX2-positive cells only. In human axioloids, NMPs support elongation by supplying new PSM and by providing a source of WNT and FGF ligands, establishing a posterior-to-anterior gradient (Yamanaka et al., 2023). This gradient positions the determination front and allows somites to form in sequence. If the pig tailbud instead supplies neural cells, this source is lost and the somitic mesoderm produced early receives no graded signal. Upon embedding in ECM, the entire somitic domain is therefore competent to segment at once. This phenotype is consistent with a study in chick embryos where non-somitic mesoderm is dorsalised by BMP inhibition, forming somites simultaneously as a “bunch of grapes”, with normal size, shape and axial identity, and without cyclic clock gene expression or a travelling wave (Dias et al., 2014). Similarly, human somitoids segment in the absence of an elongating axis: MESP2 is activated simultaneously throughout the aggregate and epithelial rosettes form without a determination front (Miao et al., 2023). These studies suggest that somite formation is a self-organising property of somitic cells and that the clock is required for the timing and spatial organisation of segmentation. As in these systems, pig TLSs showed somite boundaries but lacked ordered segmentation, consistent with tissue segmenting in the absence of a graded signal. Whether this process occurs with or without active clock oscillations remains unresolved. Analysis of clock dynamics in TLSs would address this and determine the extent to which this system resembles HES7-null human somitoids (Miao et al., 2023).

Several factors could bias pig NMPs towards neural output. In the embryo, NMPs feed both neural and mesodermal lineages (Fig. 2), so this bias is likely a property of the gastruloid system. It may also be a result of culture conditions in the induction protocol. In human axial models, dual SMAD inhibition, used to counter high baseline TGF-β signalling in human cells, represses mesodermal differentiation (Wang et al., 2026). In our protocol, the aggregation medium contained the TGF-β inhibitor SB431542, which may have a similar effect on pig NMPs. WNT dose sets the balance between SOX2 and TBXT in mouse NMPs (Gouti et al., 2014; Koch et al., 2017). Given the shifted CHIR sensitivity of pig cells, pig NMPs may need continued WNT stimulation to switch from neural to mesodermal output during elongation. However, sustaining CHIR beyond 24 h did not induce somite formation (Suppl. Fig. 6). Additionally, aggregate size could contribute to neural bias: in mouse gastruloids, small aggregates show a higher proportion of NMPs at 120 h but have depleted NMP and PSM pools by 144 h (Bennabi et al., 2025). Small aggregates are also associated with neural fate, downregulation of PSM and reduced Nodal signalling. This is consistent with reports that TGF-β inhibition represses mesodermal differentiation. We held initial cell number constant across species; however, the optimal cell number may be species-specific.

In conclusion, this study shows that somite specification and ordered segmentation can be decoupled in stem cell-based embryo models from four mammalian species. Human models, which maintain a mesoderm-producing progenitor pool, form organised somites, whereas progenitor depletion in pig models is associated with disorganised segmentation. This suggests that sustained progenitor output is a key requirement for the conversion of somitic fates into a sequentially segmenting axis. Defining the mechanisms that maintain progenitor competence and output may therefore be essential for understanding how conserved somitogenic programmes generate robust and species-specific segmentation in mammalian embryos.

## Materials and methods

### Animal ethical statement

All procedures involving animals were approved by the Animal Welfare and Ethics Review Committee (Nbr. 99) of the School of Biosciences, The University of Nottingham. The research conducted adhered to the Home Office Code of Practice guidelines for the Housing and Care of Animals used in Scientific Procedures. The use of human ES cell lines was approved by the School of Biosciences Research Ethics Committee, University of Nottingham (SBREC202223011FEO). The use of the H9 line was agreed with WiCell (Ref: 26-W0594) and HUES7 was obtained from Harvard University.

### Embryo collection

Embryos were collected from crossbred Large White and Landrace sows (2-3 years old). Each uterine horn was separated into an upper and lower half before fresh PBS+1% FCS was used to flush out porcine embryos. Uterine horns were then bisected and searched by hand for any further embryos. Embryos were stored in warm PBS+5% BSA during manipulation. Embryos were fixed in 4% PFA for 25 min at RT.

## Single-cell RNA-seq analysis

### Integration, clustering and annotation

All single-cell analyses were performed in R (v4.4.1) using Seurat (v5.4.0). Two published pig single-cell datasets were used: a gastrulation atlas spanning E11.5- E15 (Simpson et al., 2024) (GEO GSE236766) and a myogenesis dataset spanning E16-E28 (Cai et al., 2023) (GEO GSE206914). Quality control was applied as described in the original publications. Objects were split by sample, normalised, and variable features were identified, then objects were scaled and reduced by PCA. Integration features were selected across all samples (n = 2,000) and anchors identified by reciprocal PCA (dims 1:75, n.trees = 600) before integration with IntegrateData. Following integration, cells with 2,000 or fewer detected genes were removed, leaving 113,708 cells (84,412 from the gastrulation atlas and 29,296 from the myogenesis dataset). The integrated assay was scaled and reduced by PCA (75 PCs). Cells were clustered on dimensions 1:25 with SNN pruning at 1/10 and a resolution of 0.8, giving 33 clusters. To support cluster annotation, labels from the two source datasets and a mouse gastrulation atlas (Imaz-Rosshandler et al., 2024) were transferred onto the integrated cells (FindTransferAnchors/TransferData). Cluster markers were identified with FindAllMarkers (RNA assay, positive markers only, min.pct = 0.25, logfc.threshold = 0.25), and clusters were assigned identities from canonical marker expression.

### Subsetting and reannotation

Nine populations spanning the neural and skeletal muscle trajectory and neighbouring cell types were subset (43,032 cells total; 33,356 from the gastrulation atlas and 9,676 from the myogenesis dataset), reduced by PCA and reclustered (dimensions 1:30, resolution 1.8), giving 30 clusters. Cells were scored for cell type gene modules with AddModuleScore and the 30 clusters were annotated into 20 cell types from module scores and marker expression (Suppl. Fig. 1A). On closer examination of the myofibre cluster, this population was split into cardiomyocytes and skeletal myofibres using cardiac- and skeletal-specific module scores, giving 21 final cell types.

### Signalling module scores

Ligands, receptors and transcriptional targets of the WNT, FGF, NODAL, TGF- β/Activin, BMP, retinoic acid, NOTCH, YAP/TAZ and SHH pathways were grouped into modules (Suppl. Table 1) and scored with AddModuleScore across the 14 clusters covering the somitogenesis trajectory. Mean scores were scaled across clusters within each module.

## Cell culture and gastruloid generation

### Cell maintenance

Porcine (pEpiSC4, pEpiSC7), ovine (sEpiSC21) and bovine (bEpiSC99.1) epiblast stem cells were derived as described previously (Kinoshita et al., 2021). Human embryonic stem cell lines H9 and HUES7 were obtained from WiCell and Harvard University, respectively. All lines were maintained in N2B27 medium supplemented with 20 ng/ml Activin A (Peprotech), 12.5 ng/ml FGF2 (Peprotech) and 2 µM XAV939 (Tocris) (AFX medium) on Laminin-511-E8 (L511-E8; Amsbio) coated plates. Cultures were maintained at 5% CO2, 5% O2 and 38.5°C (porcine, ovine and bovine) or 37°C (human) and passaged every 3-4 days using 0.5X TrypLE.

### Gastruloid formation

Gastruloids were generated using a previously published protocol (Yamanaka et al., 2023). Cells were seeded onto laminin-coated plates and grown for 2 to 3 days with daily medium exchange, reaching ∼60% confluency. Cells were pre-treated with N2B27 supplemented with FGF2 (20 ng/ml) and CHIR99021 (between 1 and 15 µM, depending on condition) for 24 h. Cells were dissociated and reaggregated in ultra-low attachment U-bottom 96-well plates at 1000 cells per well in aggregation medium (N2B27 supplemented with FGF2 (20 ng/ml), CHIR99021 (between 0.5 and 11 µM, depending on condition), SB431542 (10 µM) and ROCK inhibitor Y27632 (10 µM)), using a 2-min centrifugation step at 250 g. 24 h after aggregation, 150 µl N2B27 was added; subsequently 150 µl medium was exchanged daily. At 48 h or 72 h after aggregation (depending on the cell line), 150 µl medium was replaced with 80 µl of 10% Matrigel (Growth-factor reduced, Corning) and cultured for a further 24 h before collection.

### 2D CHIR dose-response experiment

To assess species-specific CHIR sensitivity, pEpiSC7 and H9 cells were seeded in a 96-well plate at 2,500 cells per well on laminin-coated plates and cultured for 2 days in AFX medium. Cells were then subjected to media changes as per the gastruloid formation protocol, without the aggregation step. Briefly, cells were pre-treated for 24 h with N2B27 supplemented with FGF2 (20 ng/ml) and CHIR99021 (1, 3, 5, 7, 11 or 15 µM), followed by a further 24 h in aggregation medium (N2B27 with FGF2 (20 ng/ml), SB431542 (10 µM) and CHIR99021 (0.5, 1, 3, 5, 9 or 11 µM)), giving a 6x6 matrix of CHIR combinations per cell line. Cells were fixed 48 h after the start of pre-treatment.

## Immunostaining

### Gastruloids and whole-mount embryos

Samples were fixed for 25 min in 4% paraformaldehyde (PFA) at room temperature (RT) and stored in PBS containing 0.05% sodium azide at 4°C. Prior to immunostaining, samples were washed twice in PBS with 0.1% Triton-X. Embryos were cut into parts prior to staining. Samples were permeabilised and blocked for 3 hours at RT in PB buffer (PBS with 5% donkey serum, 5% BSA, and 1% Triton-X). Samples were then incubated in primary antibodies diluted in PB buffer overnight at 4°C and incubated in secondary antibodies with Hoechst (1:1000, Thermo Fisher H3570) for 2 hours at RT in PB buffer. Following both antibody incubations, samples were washed four times for 15 min each in PBS containing 0.1% Triton-X. After immunostaining, samples were incubated overnight at 4 °C in ScaleS4 clearing solution (Hama et al., 2015). Cleared samples were mounted on cavity slides in Vectashield mounting medium before imaging. Antibodies used are listed in Suppl. Table 2.

### Cells in monolayer

Cells in well plates were fixed and immunostained as described for gastruloids, with reduced incubation times for fixation (15 min), blocking/permeabilisation (1 h), secondary antibody incubation (1 h) and washes (4 x 5 min).

## Imaging

### Confocal imaging

Whole-mount embryos and gastruloids were imaged across multiple z-planes using a BC43 confocal microscope (Oxford Instruments), with 10x, 20x or 40x objectives, with z-intervals of 0.6 or 1 µm for most images, varying between panels. Images are displayed as single z-planes or maximum-intensity projections. Tiled acquisitions were stitched and, where appropriate, deconvolved using Imaris. For display, the TBX6 channel in Figs 3E and 6C was background corrected using a Gaussian filter (σ = 300 px), morphological opening (radius 5 px) and a median filter (kernel size 5 px). In Fig. 3D, speckle in the pig MEOX1 channel was removed by greyscale morphological opening (disk, radius 4 px); the human MEOX1 channel was not filtered. These corrections were applied for display only.

### Widefield imaging

For morphometric analyses, fixed gastruloids were stained with Hoechst alone, transferred individually to U-bottom 96-well plates and imaged over a short z-stack (10 µm spacing) at 10x magnification using a Zeiss Axiovert. Masks are displayed in Suppl. Fig. 2A. Cells stained in 2D were imaged at 20x magnification (single plane) on the same system.

### Brightfield imaging

Brightfield images of gastruloids were acquired on a Leica Mateo TL at 10x magnification. Live imaging was performed on a BC43 confocal microscope (Oxford Instruments) fitted with an incubation chamber to maintain the appropriate temperature, humidity, O2 and CO2 levels for each cell line, with brightfield z-stacks (10/16 slices) acquired every 30 min for 20 h from the time of MG embedding, at 10x magnification.

## Quantitative image analysis

### Gastruloid segmentation and morphometrics

Image segmentation and quantification were performed in Python. Gastruloid masks were obtained from two different image types. Human and pig gastruloids (Fig. 3C, Suppl. Fig. 2A, B) were segmented from maximum-intensity projections of Hoechst-stained gastruloids using Cellpose (Stringer and Pachitariu, 2025) with the estimated diameter adjusted for the smaller size of the gastruloids at 24 h. Where more than one object was returned, the object closest in area to the median for that condition was retained. Cow and sheep gastruloids (Fig. 5C, Suppl. Fig. 4) were segmented from brightfield images using the Segment Anything Model (SAM) (Kirillov et al., 2023). All masks were inspected visually and corrected or excluded where segmentation had failed. Area and aspect ratio were extracted from each mask, with aspect ratio calculated as the ratio of major to minor axis length.

### Live imaging analysis and somite boundary detection

Gastruloid masks were obtained from segmentation of time-lapse images (40 timepoints, 30 min intervals) using SAM 2.1 in video mode. Midline tracing for each frame was adapted to the morphology of each species. For human, the mask was skeletonised (scikit-image) and the midline was defined as the longest path through the skeleton, extended to the mask boundary. For pig, the midline was the straight line along the first principal component of the mask pixel coordinates, passing through the mask centroid. Boundary positions were annotated manually. Images from all replicates of both species (n = 5) and all timepoints (400 frames) were pooled and presented in randomised order, blinded to species, sample and timepoint. Each boundary position was then projected onto the nearest point of the midline of its frame and assigned a position, defined in µm as the length from the anterior end.

### Nuclear segmentation and marker quantification

For the monolayer CHIR dose-response experiment, nuclei were segmented from the Hoechst channel using Cellpose (nuclei_cp3), following unsharp-mask pre-processing to enhance nuclear boundaries. For each channel, background was estimated as the median intensity of non-nuclear pixels and subtracted, and mean intensity extracted per nucleus. Intensities were log₁₀-transformed and a two-component Gaussian mixture model fitted to the pooled distribution for each marker and cell line, with the threshold set at the crossover point between components. The percentage of positive cells was calculated per condition and averaged across 3 images per well per experiment and across 3 independent experiments. Conditions with an average cell number per well below 50% of the median across all conditions were excluded from quantification. For 3D gastruloid images, nuclei were segmented from the Hoechst channel using StarDist (Schmidt et al., 2018), applied to each z- slice following unsharp-mask pre-processing. 3D nuclei were then reconstituted by linking masks together across consecutive slices by pixel overlap. Background was subtracted per channel, either as the median intensity of non-nuclear voxels (Fig. 6E) or as a Gaussian-blurred background surface subtracted per slice followed by a median filter (Fig. 3F). Mean intensity was extracted per nucleus for each marker channel. Marker intensity distributions in 3D samples were not consistently bimodal, reflecting variable antibody and light penetration within samples, so the thresholding strategy was adapted to each experiment. For Fig. 3F, thresholds were set per marker as fixed percentiles of the intensity distribution pooled across all nuclei. For Fig. 6E, a two-component Gaussian mixture model was fitted to the log₁₀- transformed intensities at each timepoint for each marker and cell line; the timepoint showing the cleanest bimodal separation was used as the threshold source and applied across all timepoints for that cell line.

### Statistics

Replicates for each experiment are indicated in the figure legends. Statistical analyses were performed in R. Live-imaging measurements (Fig. 6B) were compared between species using a two-tailed Mann-Whitney U test. Marker quantification (Fig. 6E) was analysed by two-way ANOVA with Šídák’s multiple comparisons test.

### Code and data availability

The single-cell RNA-seq data analysed in this study are publicly available. The pig gastrulation atlas (Simpson et al., 2024) is available from the NCBI Gene Expression Omnibus (GEO) under accession GSE236766, and the pig myogenesis dataset (Cai et al., 2023) under accession GSE206914. The integrated and annotated Seurat objects generated in this study are available on Zenodo (https://doi.org/10.5281/zenodo.23835905). Code for integration, clustering, annotation and module scoring is available on GitHub (https://github.com/AlberioLab/pig-somitogenesis-scrnaseq).

## Declaration of AI use

During the preparation of this work, the authors used AI to assist with writing the code for the single-cell integration and annotation pipeline, for the image analysis pipelines, and language editing.

## Author contributions

CK, LS and RA designed experiments and wrote the paper. CK performed experiments. LS contributed to the scRNA-seq analysis. CA provided the HUES7 cell line. CLRM discussed experiments and results. RA supervised the project.

## Acknowledgements

We thank members of the Alberio lab for helpful discussions and support throughout the project and Osvaldo Chara for valuable suggestions.

## Supplementary information

**Supplementary Figure 1.**
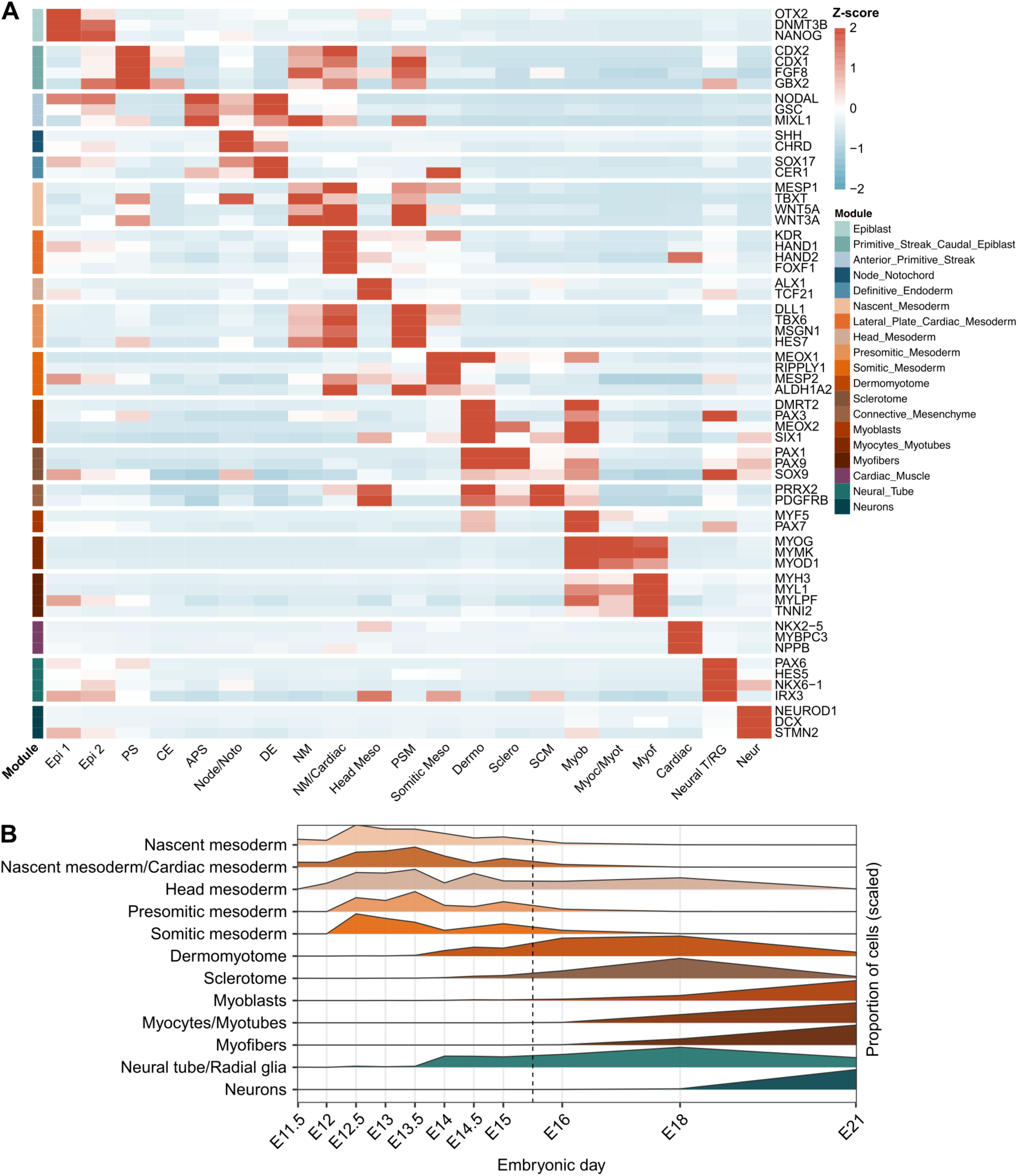
Marker gene expression and population dynamics of annotated cell types. (A) Heat map of marker gene expression across the annotated clusters shown in Fig. 1A. Rows are marker genes, grouped into cell type modules (colour bar, left) and labelled on the right. Columns are clusters. Colour shows the aggregated expression of each gene per cluster (z-score across clusters; RNA assay). (B) Relative abundance of selected populations across embryonic days (E11.5-E21). Cell proportions were averaged across samples of each stage and scaled to each population’s maximum. The dashed line marks the boundary between the two datasets. Epi, epiblast; PS, primitive streak; CE, caudal epiblast; APS, anterior primitive streak; Node/Noto, node/notochord; DE, definitive endoderm; NM, nascent mesoderm; NM/Cardiac, nascent mesoderm/cardiac mesoderm; Meso, mesoderm; PSM, presomitic mesoderm; Dermo, dermomyotome; Sclero, sclerotome; SCM, somitic connective mesenchyme; Myob, myoblasts; Myoc/Myot, myocytes/myotubes; Myof, myofibres; Neural T/RG, neural tube/radial glia; Neur, neurons.

**Supplementary Figure 2.**
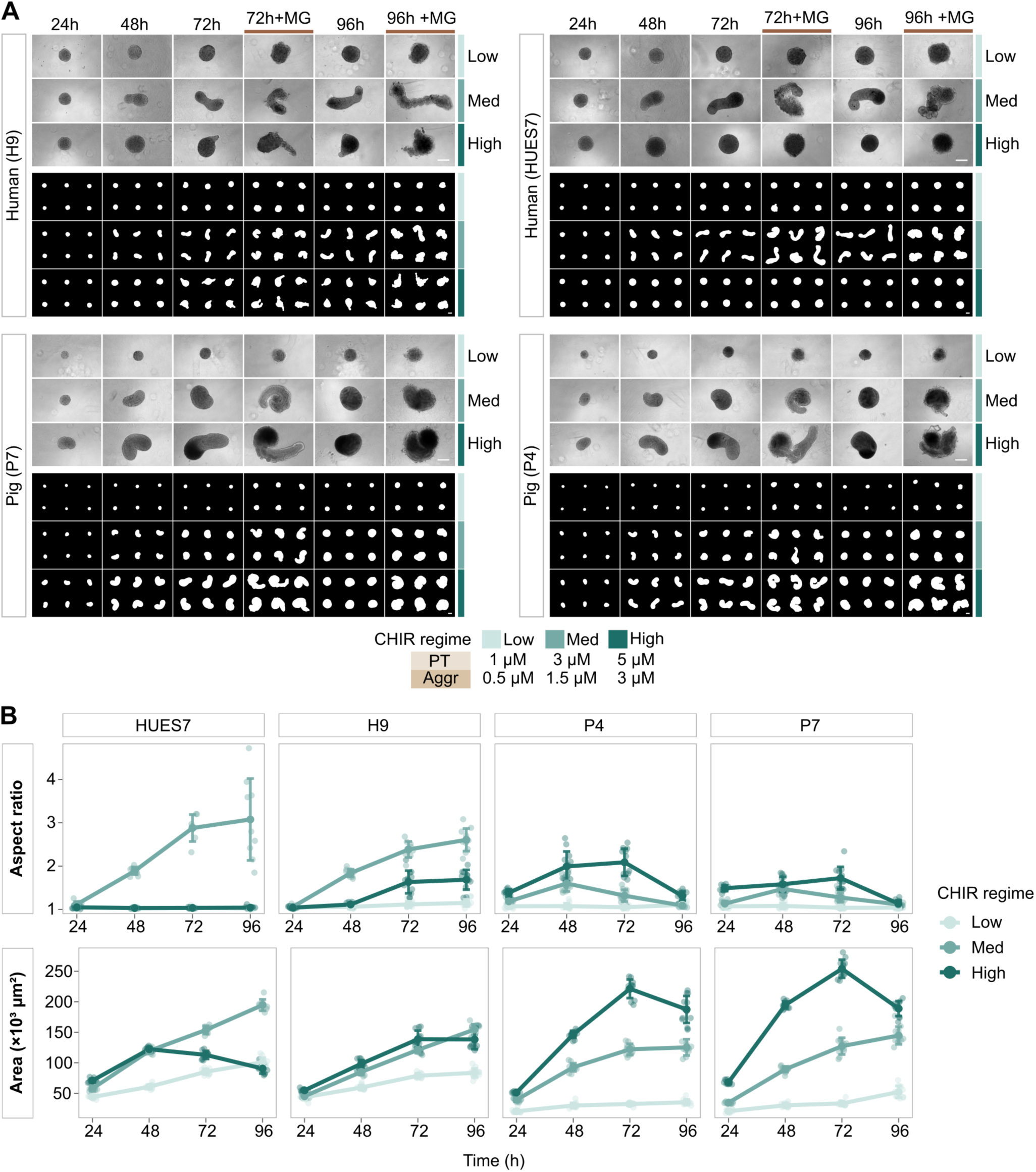
Morphological characterisation of human and pig gastruloids from two different cell lines per species. (A) Brightfield images of human (HUES7 and H9) and pig (P4 and P7) gastruloids from 24 to 96 h under low, med and high CHIR regimes. 72 h + MG and 96 h + MG indicate gastruloids embedded in MG 24 h before imaging. Binary masks obtained from gastruloid segmentation (six per condition) are displayed beneath each brightfield panel. Scale bars, 200 µm. (B) Aspect ratio and area of human (HUES7 and H9) and pig (P4 and P7) gastruloids over time under three CHIR regimes. Data show mean ± s.d., with points representing individual gastruloids (n = 7-12 gastruloids per condition).

**Supplementary Figure 3.**
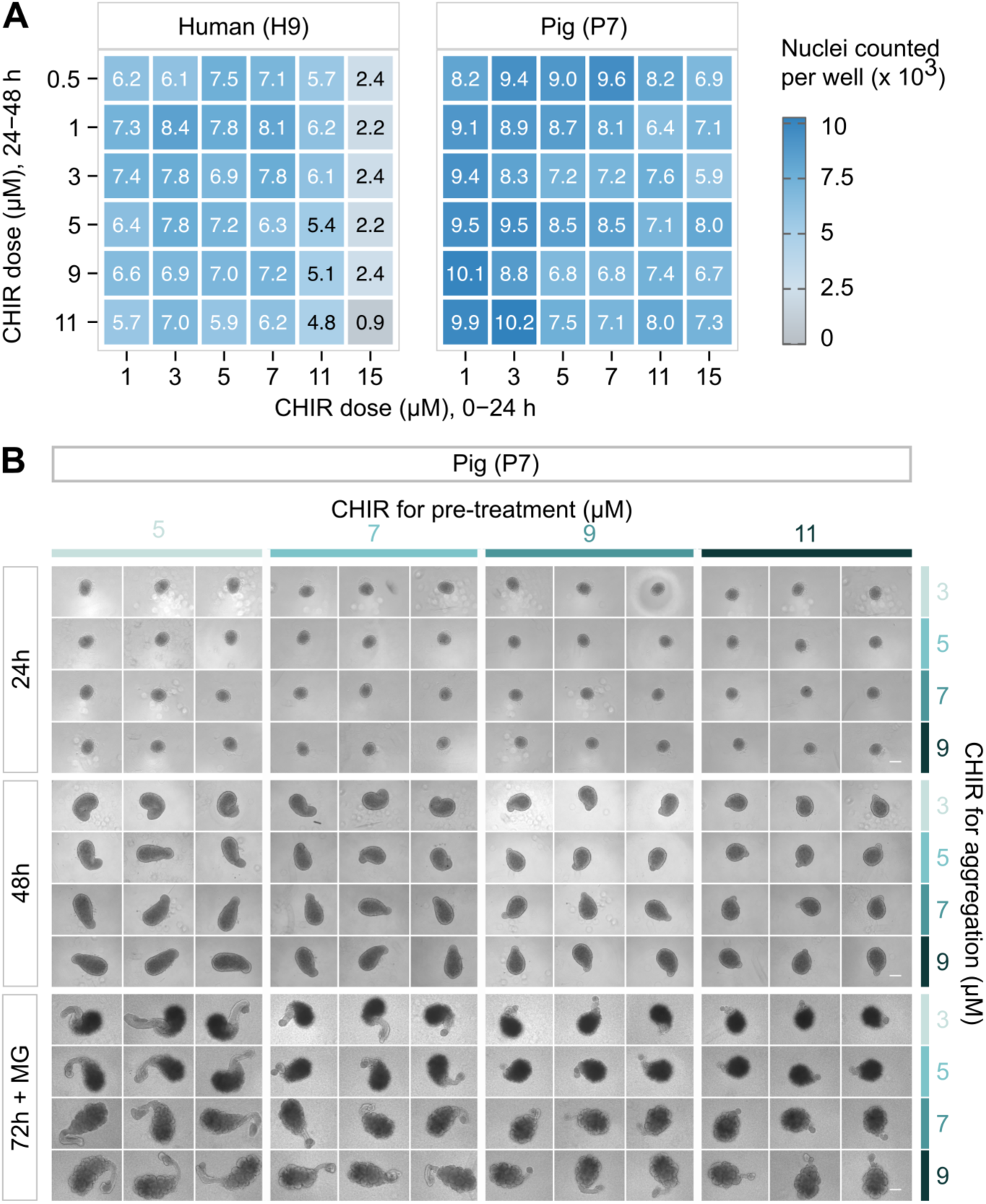
Cell counts in 2D CHIR dose-response experiment and pig gastruloid morphology across CHIR ranges. (A) Average cell count per well in human (H9) and pig (P7) monolayer experiment across the CHIR dose matrix shown in Fig. 4B. The x-axis shows the CHIR dose at 0-24 h and the y-axis the dose at 24-48 h. Each value is the mean of 3 regions per well across 3 independent experiments. (B) Brightfield images of pig (P7) gastruloids at 24, 48 and 72 h + MG across a matrix of CHIR concentrations at pre-treatment (columns) and aggregation (rows). Three replicates are shown per condition. Scale bar, 200 µm.

**Supplementary Figure 4.**
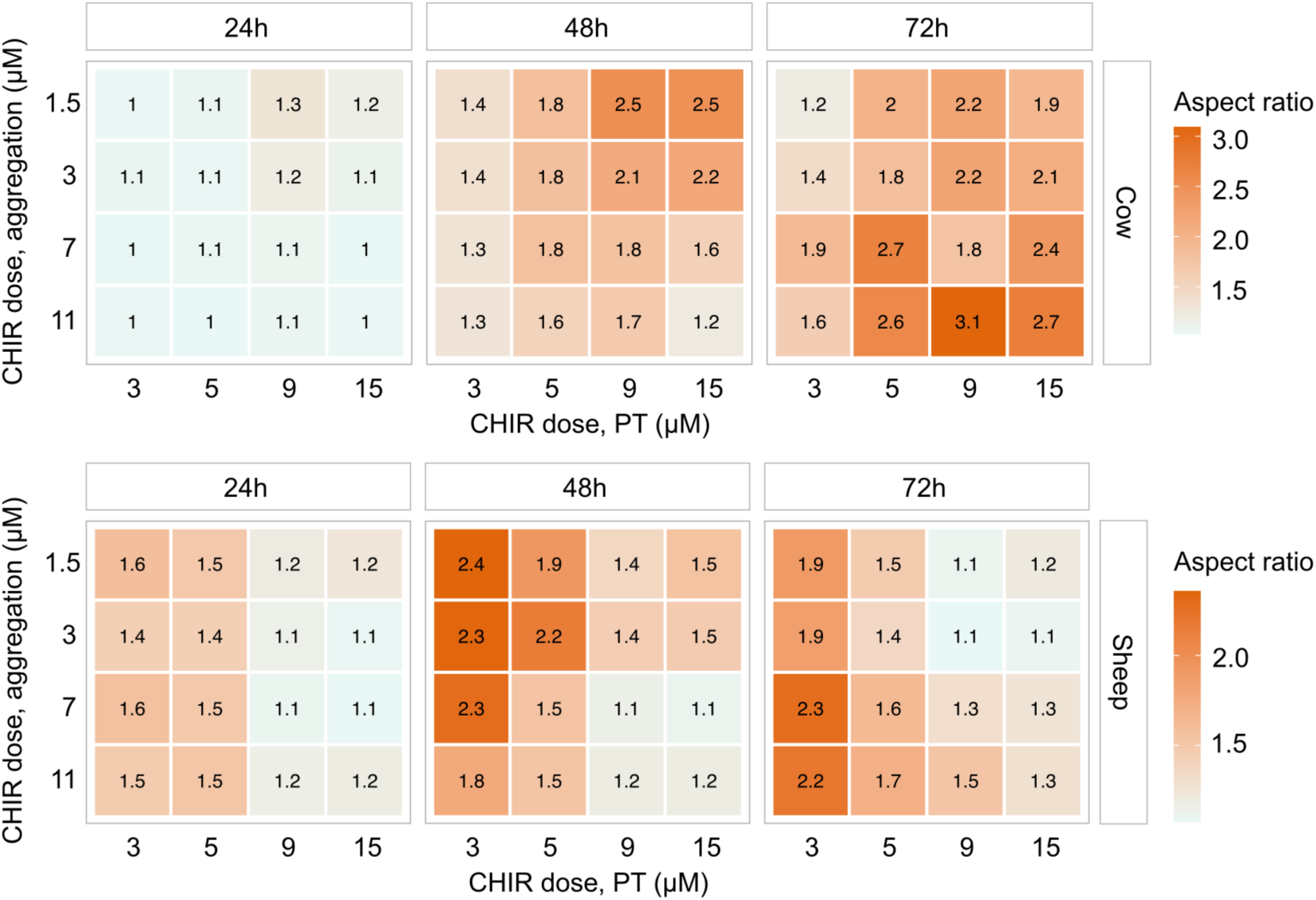
Aspect ratio of cow and sheep gastruloids across a range of CHIR regimes. Aspect ratio of cow (top) and sheep (bottom) gastruloids at 24, 48, and 72 h across a matrix of CHIR concentrations at pre-treatment and aggregation. Values are the mean aspect ratio per condition (n = 6 gastruloids per condition).

**Supplementary Figure 5.**
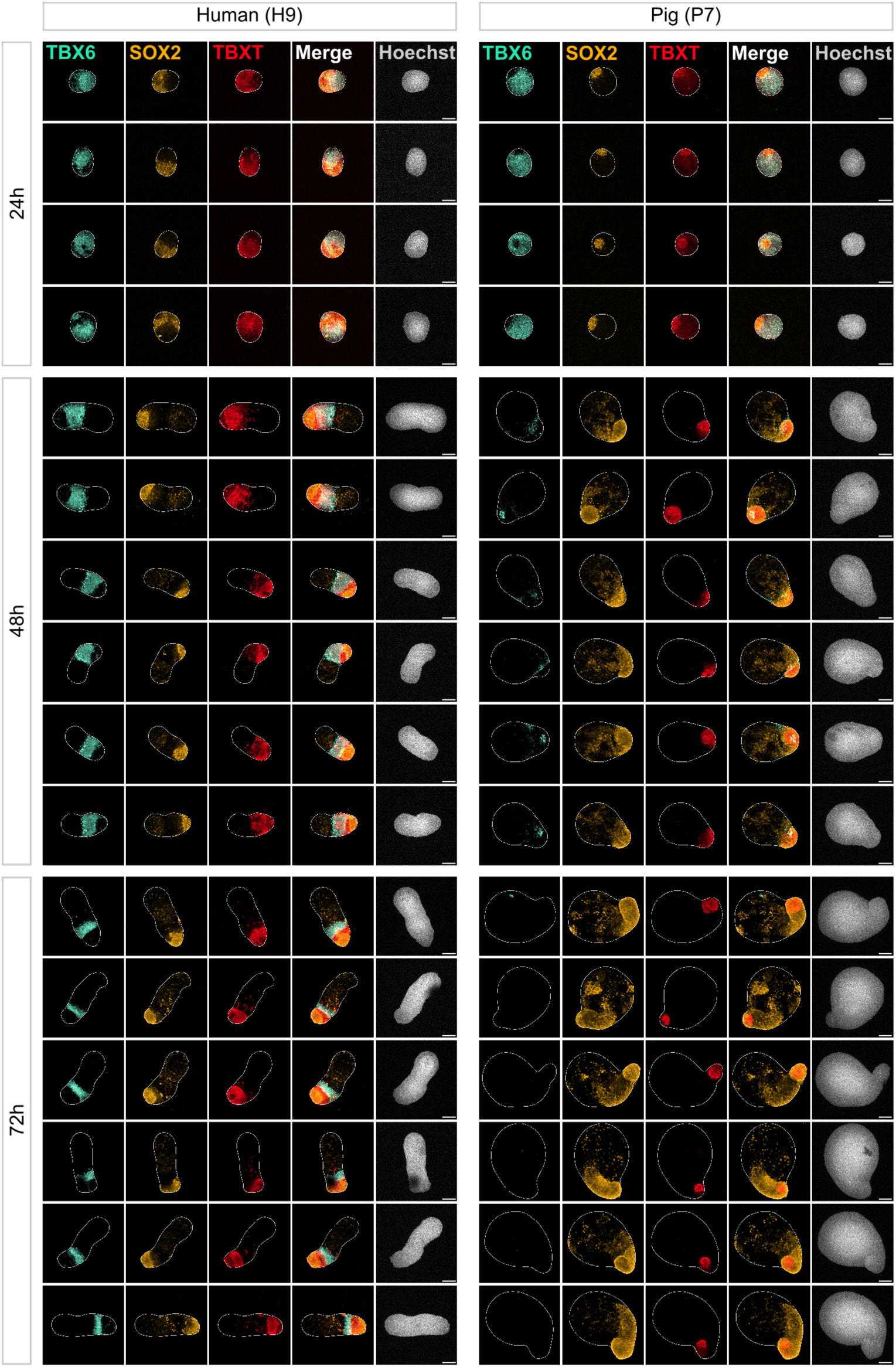
Extended immunofluorescence data of human and pig gastruloids at 24, 48 and 72. **h** Immunofluorescence images (maximum-intensity projections) of human (H9) and pig (P7) gastruloids (CHIR pre-treatment/aggregation: 3/1.5 µM for human, 11/9 µM for pig) at 24, 48 and 72 h stained for TBX6, SOX2 and TBXT, with Hoechst. Each row shows a replicate gastruloid (n = 4 gastruloids at 24 h and n = 6 at 48 and 72 h per species). White lines indicate the gastruloid outline. Scale bars, 200 µm.

**Supplementary Figure 6.**
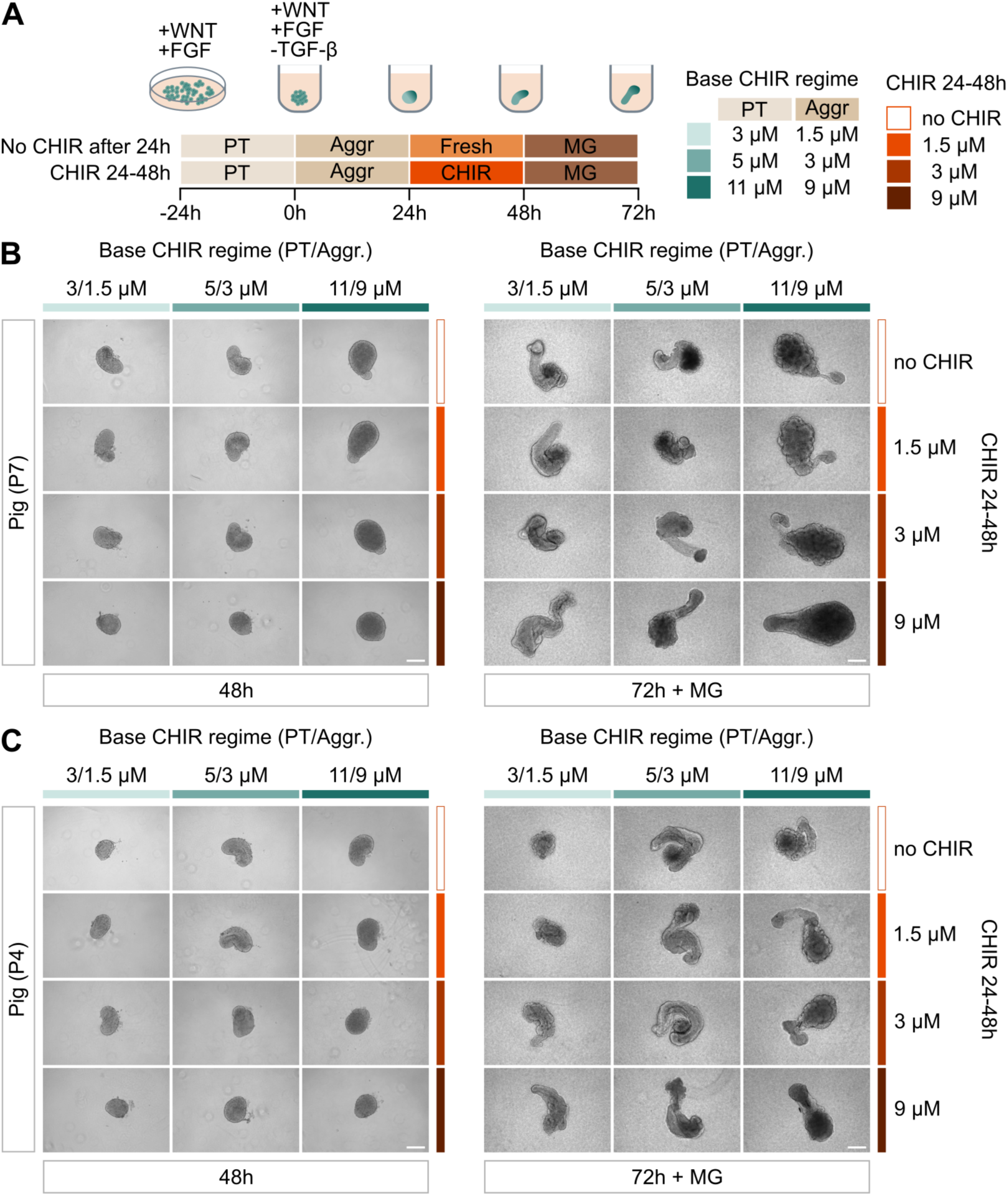
Effect of additional CHIR treatment between 24 and 48 h on pig gastruloid morphology. (A) Schematic of the protocol. Gastruloids were generated under three base CHIR regimes (pre-treatment/aggregation: 3/1.5, 5/3 and 11/9 µM). In control gastruloids, medium was replaced with fresh medium at 24 h (“Fresh”). In the CHIR 24-48 h condition, medium was replaced with medium containing 1.5, 3 or 9 µM CHIR at 24 h, and gastruloids were embedded in MG at 48 h. Time is shown as hours from aggregation. (B) Brightfield images of pig (P7) gastruloids at 48 h and 72 h + MG under the indicated base CHIR regimes (columns) and CHIR concentrations from 24 to 48 h (rows). (C) Same as B, for pig (P4) gastruloids. One gastruloid per condition is shown. Scale bars, 200 µm.

**Supplementary Movie 1. Live imaging of a human axioloid.**

Brightfield time-lapse of a human (H9) axioloid recorded over 20 h, starting at MG embedding (0 h; 72 h of the protocol). Images were acquired every 30 min and are displayed as maximum-intensity projections of 10 z-slices. Time stamps indicate hours since MG embedding. Scale bar, 200 µm.

**Supplementary Movie 2. Live imaging of a pig TLS.**

Brightfield time-lapse of a pig (P7) TLS recorded over 20 h, starting at MG embedding (0 h; 48 h of the protocol). Images were acquired every 30 min and are displayed as maximum-intensity projections of 16 z-slices. Time stamps indicate hours since MG embedding. Scale bar, 200 µm.

**Supplementary Table 1.** Genes used for signalling pathway module scores.

| **WNT** | **FGF** | **NODAL** | **TGF-β/Activin** | **BMP** | **Retinoic acid** | **NOTCH** | **YAP/TAZ** | **SHH** |
|---|---|---|---|---|---|---|---|---|
| *AXIN2* | *FGF8* | *NODAL* | *SERPINE1* | *ID1* | *ALDH1A2* | *DLL1* | *CCN1* | *GLI1* |
| *NKD1* | *DUSP6* | *GDF1* | *SKIL* | *ID2* | *RDH10* | *DLL3* | *CCN2* | *PTCH1* |
| *SP5* | *SPRY2* | *GDF3* | *INHBA* | *ID3* | *STRA6* | *HES7* | *ANKRD1* | *HHIP* |
| *NOTUM* | *SPRY4* | *TDGF1* | *INHBB* | *MSX1* | *CRABP2* | *HES1* | *AMOTL2* | *SHH* |
| *LEF1* | *ETV4* | *FOXH1* | *TGFB1* | *MSX2* | *DHRS3* | *LFNG* | *THBS1* | *GLI2* |
| *TCF7* | *ETV5* | *PITX2* | *TGFB2* | *BMP2* | *RARB* | *HEY1* | *AXL* | *SMO* |
| *TCF7L2* | *FGFR1* | *LEFTY1* | *TGFB3* | *BMP4* | *ALDH1A1* | *HEY2* | *CRIM1* | *PTCH2* |
| *WNT3* | *FGFR2* | *LEFTY2* | *TGFBR1* | *BMPR1A* | *ALDH1A3* | *NRARP* |  | *GAS1* |
| *WNT3A* | *DUSP4* | *ACVR1B* | *TGFBR2* | *BMPR2* | *RARA* | *NOTCH1* |  | *CDON* |
| *WNT8A* | *DUSP5* | *SMAD2* | *SMAD2* | *SMAD1* | *RARG* | *NOTCH2* |  | *BOC* |
| *WNT5A* | *SPRY1* | *SMAD3* | *SMAD3* | *SMAD5* | *RXRA* | *JAG1* |  | *DISP1* |
| *ROR2* | *SPRED1* | *SMAD7* | *SMAD7* | *SMAD6* | *RXRG* | *RBPJ* |  | *SCUBE2* |
| *LRP6* | *SPRED2* | *ACVR2A* | *ACVR2A* | *BAMBI* | *CRABP1* | *HEYL* |  | *HHAT* |
| *RNF43* | *IL17RD* | *ACVR2B* | *ACVR2B* | *ID4* | *RBP1* | *DTX1* |  | *SUFU* |
| *ZNRF3* | *ETV1* | *ACVR1C* | *GDF11* | *BMP5* | *FGF18* | *MFNG* |  | *GPR161* |
| *APCDD1* | *FGF4* |  | *LTBP1* | *BMP6* |  | *RFNG* |  |  |
| *FZD5* | *FGF17* |  |  | *BMP7* |  | *MIB1* |  |  |
| *FZD7* | *CYP26A1* |  |  | *ACVR1* |  |  |  |  |
| *FZD8* | *FLRT3* |  |  | *BMPR1B* |  |  |  |  |
|  |  |  |  | *SMAD9* |  |  |  |  |
|  |  |  |  | *GDF6* |  |  |  |  |

**Supplementary Table 2.** List of antibodies.

| **Antibody** | **Reference** | **Dilution** |
|---|---|---|
| Primary antibodies |  |  |
| Anti-TBXT (Goat) | R&D Systems Cat# AF2085, RRID:AB_2200235 | 1:500 |
| Anti-TBX6 (Rabbit) | Abcam Cat# ab38883, RRID:AB_778274 | 1:250 |
| Anti-SOX2 (Rat) | Thermo Fisher Scientific Cat# 14-9811-82, RRID:AB_11219471 | 1:1500 |
| Anti-SOX1 (Rabbit) | Cell Signaling Technology Cat# 4194, RRID:AB_1904140 | 1:200 |
| Anti-PAX3 (Rabbit) | Thermo Fisher Scientific Cat# 38-1801, RRID:AB_2533359 | 1:250 |
| Anti-MEOX1 (Rabbit) | Atlas Antibodies Cat# HPA045214, RRID:AB_10959299 | 1:700 |
| Secondary antibodies |  |  |
| Donkey anti-rat IgG, Alexa Fluor 555 | Thermo Fisher Scientific Cat# A78945, RRID:AB_2910652 | 1:1000 |
| Donkey anti-goat IgG, Alexa Fluor 680 | Thermo Fisher Scientific Cat# A-21084, RRID:AB_2535741 | 1:1000 |
| Donkey anti-rabbit IgG, Alexa Fluor 488 | Thermo Fisher Scientific Cat# A-21206, RRID:AB_2535792 | 1:1000 |

